# Super-silencers regulated by chromatin interactions control apoptotic genes

**DOI:** 10.1101/2022.01.17.476559

**Authors:** Ying Zhang, Kaijing Chen, Yichao Cai, Akiko Nambu, Yi Xiang See, Chaoyu Fu, Anandhkumar Raju, Manikandan Lakshmanan, Motomi Osato, Vinay Tergaonkar, Melissa Jane Fullwood

## Abstract

Human silencers have been shown to exist and regulate developmental gene expression. However, the functional importance of human silencers needs to be elucidated such as the working mechanism and whether they can form “super-silencers”. Here, through interrogating two putative silencer components of *FGF18* gene, we found that two silencers can cooperate via compensated chromatin interactions to form a “super-silencer”. Furthermore, double knock-out of two silencers exhibited synergistic upregulation of *FGF18* expression and changes of cell identity. To disturb the “super-silencers”, we applied combinational treatment of an EZH2 inhibitor GSK343, and a REST inhibitor, X5050 (“GR”). We found that GR led to severe loss of TADs and loops, while the use of just one inhibitor by itself only showed mild changes. Such changes of TADs and loops may due to reduced CTCF protein level observed upon GR treatment. Moreover, GSK343 and X5050 worked together synergistically to upregulate the apoptotic genes controlled by super-silencers, and thus gave rise to antitumor effects including apoptosis, cell cycle arrest and tumor growth inhibition. Overall, our data demonstrated the first example of a “super-silencer” and showed that combinational usage of GSK343 and X5050 could potentially lead to cancer ablation through disruption of “super-silencers”.

## Introduction

Cis-regulatory elements are important for controlling gene expression including active elements such enhancers and super-enhancers (SEs) (Pott & Lieb, 2015; Whyte et al., 2013) and repressive elements such as silencers (Cai et al., 2021; Doni Jayavelu, Jajodia, Mishra, & Hawkins, 2020; D. Huang, Petrykowska, Miller, Elnitski, & Ovcharenko, 2019; Ngan et al., 2020). Enhancers are enriched with H3K4me1 and H3K27ac (Visel et al., 2009) and there are multiple enhancers that can control the same gene’s expression (Hong, Hendrix, & Levine, 2008; Moorthy et al., 2017; Werner, Hammer, Wahlbuhl, Bosl, & Wegner, 2007). SEs are stitched by multiple enhancers and bound by master transcription factors such as OCT4 and SOX2 to drive expression of genes that define cell identity (Whyte et al, 2013).

Different components of SE can work in different modes to activate target gene expression: hierarchical (J. Huang et al., 2018; Shin et al., 2016), additive (Hay et al., 2016; Moorthy et al., 2017), and redundant (Frankel et al., 2010; Osterwalder et al., 2018). For example, Shin et al (Shin et al., 2016) dissected the STAT5-driven Wap SE and found hierarchy among enhancers. Specifically, they found that constituents of super-enhancers formed in temporal order which showed that earliest formed constituent is essential for subsequent constituents to be established and the latter constituents have the highest level of regulatory activity. Similarly, Hay et al (Hay et al., 2016) interrogated the α-globin SE *in vivo*. However, they found that each constituents contributes individually in an additive manner and not in synergy. As for the redundant theory, there are multiple reports (Frankel et al., 2010; Osterwalder et al., 2018) which demonstrated that enhancers control genes in a spatio-temporal pattern. Enhancers might be redundant in one environment but functional in a more sensitized environment, therefore they contribute to the phenotype robustness when facing environmental and genetic variability.

SEs have been shown to be acquired by the key oncogenes such as the *MYC* oncogene (Hnisz et al., 2013) in cancer to drive the process of tumorigenesis. SEs are highly associated with chromatin interactions (Cao et al., 2017) and they can connect with the target oncogene via long-range chromatin interactions (Babu & Fullwood, 2015; Bradner, Hnisz, & Young, 2017). Altered chromatin interactions have been observed to drive the expression of oncogenes such as *TERT* (Akincilar et al., 2016). Therefore, inhibition of cancer specific SEs became a new direction to investigate for cancer therapies (Bradner et al., 2017). Pharmacological inhibition of transcriptional activators that are involved in SE function can be one strategy to treat cancer (Loven et al., 2013). For example, JQ1 and trametinib, as BET inhibitors, are now in clinical trial and show the potential to treat colorectal cancer (Y. Ma et al., 2017) through targeting BET family proteins especially BRD4 to disrupt the SEs (Loven et al., 2013). Another strategy to disrupt the SEs maybe disruption of long range chromatin interactions between SEs and target oncogenes. Recently, one class of anti-cancer drugs, curaxins (Gasparian et al., 2011) have been found to disrupt long range chromatin interactions both *in vitro* and *in vivo*, and thus supressing the expression of key oncogenes especially *MYC* family genes (Kantidze et al., 2019).

Silencers, which are regions of the genome that are capable of repressing gene expression, can be predicted by various methods based on histone marks and chromatin interactions (Cai et al., 2021; Doni Jayavelu et al., 2020; D. Huang et al., 2019; Ngan et al., 2020). Recently, different methods have been proposed to identify silencers in human and mouse including correlation between H3K27me3-DNaseI hypersensitive site and gene expression (D. Huang et al., 2019), subtractive approach (Doni Jayavelu, Jajodia, Mishra, & Hawkins, 2020) and PRC2 Chromatin Interaction Analysis with Paired-End Tag sequencing (ChIA-PET) (Ngan et al., 2020). Our previous work also demonstrated a method to identify silencers through ranking and stitching the histone H3K27me3 peaks, through which to identify H3K27me3-rich regions (MRRs) as human silencers (Cai et al., 2021). Moreover, we validated the existence of silencers and showed that they can control cell identity related genes (Cai et al., 2021), which was also agreed with several other studies (Cai et al., 2021; Doni Jayavelu et al., 2020; D. Huang et al., 2019; Ngan et al., 2020). This results indicated the cell type specificity of silencers, which was similar to SEs. Similar to SEs, we also found that MRRs were highly associated with chromatin interactions and indeed, we validated two looping silencers. The above are consistent with the characteristics of SEs, which showed that MRRs exhibit cell type specificity and enrichment for chromatin interactions. Therefore, we speculated that MRRs could be “super-silencers”. Here we define “super-silencer” as a genomic region of high H3K27me3 signal comprising multiple silencers that can work as a whole entity to repress transcription of genes involved in cell identity.

Studies about human silencers are still in the early stages and the functional importance of silencers has not been validated yet. Studies of enhancers have revealed their impact on specificity and robustness of transcriptional regulation and their importance during development and evolution (Hnisz et al., 2013; Long, Prescott, & Wysocka, 2016; Pennacchio, Bickmore, Dean, Nobrega, & Bejerano, 2013; Whyte et al., 2013). Moreover, epigenetic drugs that target SEs have shown good efficiency in terms of killing cancer cells (Loven et al., 2013). Similarly, it is crucial to address some fundamental questions about silencers especially validating the putative silencers and exploring their working mode to demonstrate whether they can be “super-silencers”. Moreover, similar to drugs that target SEs, finding drugs that can target silencers or “super-silencers” may reveal another dimension for cancer therapies.

Here, we demonstrated that two silencer components inside of one MRR can cooperate as a “super-silencer” through compensated chromatin interactions to repress *FGF18* gene. Double knock-out (DKO) of these two silencer components caused synergistic upregulation of *FGF18* gene expression and synergistic cell identity changes as well. Through Hi-C and 4C, we found that the 3D genome reorganization that accompanies these epigenomic changes underlies such synergism. We disrupted the super-silencers and chromatin interactions with the epigenetic drugs GSK343 and X5050 targeting EZH2 and REST respectively. Surprisingly, we found that the single treatment of either GSK343 or X5050 mildly changed the genome organization and cancer cell viability, while combinational treatment of GSK343 and X5050 exerted synergistic effects. Specifically, we revealed that combinational treatment of GSK343 and X5050 can upregulate apoptotic genes controlled by super-silencers, which may explain the synergistic antitumor effect that we observed. Combinational treatment of GSK343 and X5050 led to reduced CTCF protein levels, which could explain the large disruption of TADs and loops. Taken together, our results suggest that the usage of combinational treatment to target super-silencers and 3D genome organization should be further researched as a potential future cancer therapy.

## Results

### Removal of two silencers in the same MRR leads to synergistic upregulation of *FGF18* expression and growth inhibition

In our previous work, we experimentally validated two powerful distal silencers (Silencers loop to *IGF2* gene and *FGF18* gene respectively) (Cai et al., 2021). To investigate the working mode of silencers, we further dissected the two component silencers within the MRR distal to the *FGF18* gene (silencer1, “S1” and silencer2, “S2”). We generated individually or combinatorial (double) CRISPR knock-out (KO) clones (S1KO, S2KO and DKO) (Figure 1A, Figure S1A-B).

**Figure 1.**
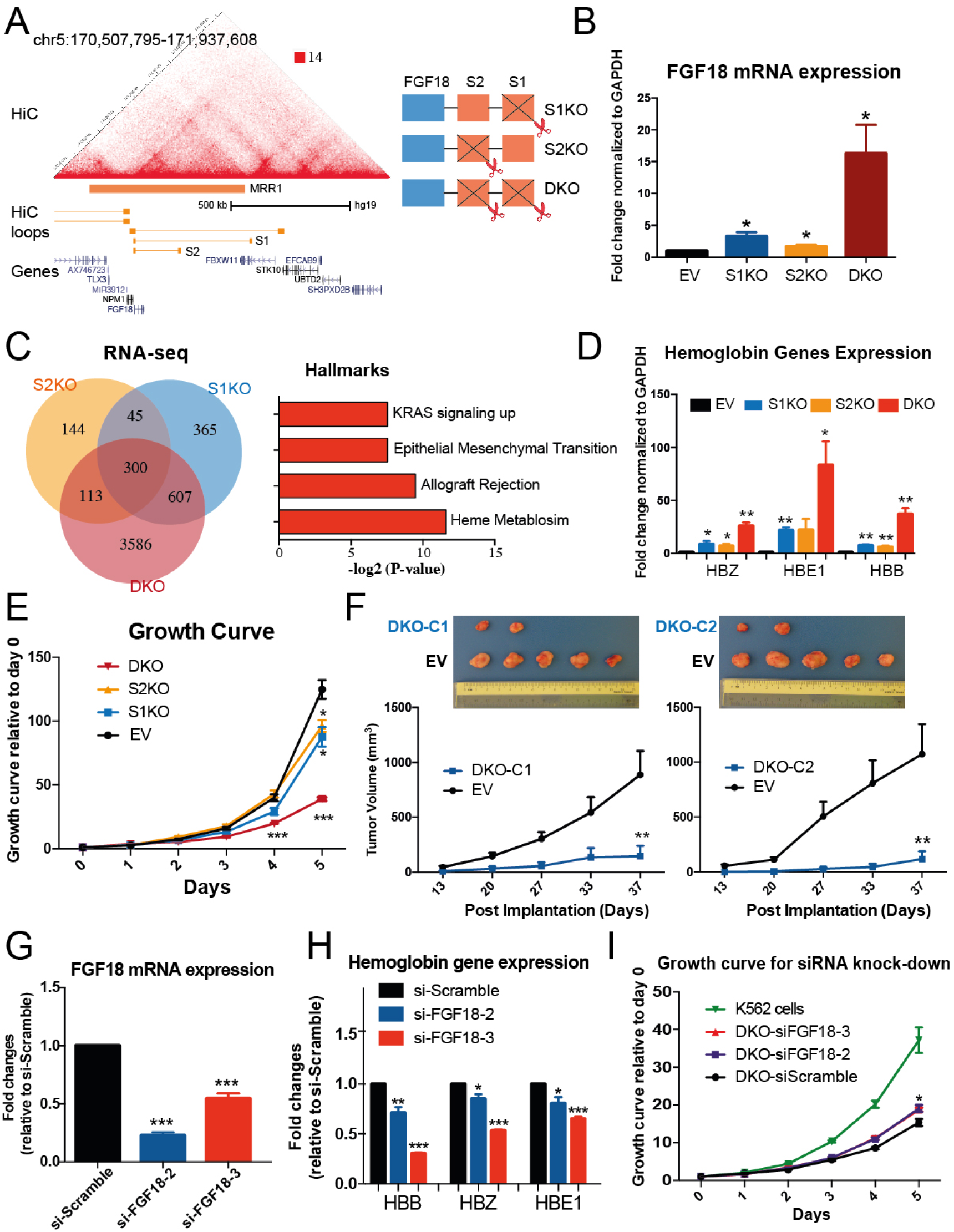
Two silencers removal led to synergistic upregulation of *FGF18* expression and growth inhibition. **A**. Hi-C matrix and loops (Rao et al., 2014) at the MRR1 region shows two looing silencers (S1 and S2) have chromatin interaction to *FGF18* gene. A schematic demonstrated the strategy to CRISPR out different silencer components to generate S1 knock-out cells (S1KO), S2 knock-out cells (S2KO) and double knock-outs of S1 and S2 (DKO). **B**. RT-qPCR of FGF18 gene expression in the vector control clone (“Empty Vector”; “EV”), S1KO cells, S2KO cells and DKO cells. Fold change was plotted normalized to GAPDH. **C**. Venn diagram showed the significant differentially expressed genes in the RNA-seq data in S1KO, S2KO and DKO cells as compared to EV cells. Gene Set Enrichment Analysis (GSEA) was performed using the intersection of significant differentially expressed genes in different RNA-seq (300 genes) against cancer hallmark database (Subramanian et al., 2005). Data was shown as –log_2_(p value). **D**. RT-qPCR of hemoglobin genes (*HBZ*, *HBE1* and *HBB*) expression in the EV cells, S1KO cells, S2KO cells and DKO cells. Fold change was plotted normalized to GAPDH. **E**. Growth curve in EV cells and different knock-out cells. Data was calculated as the fold change against day 0. **F**. Tumor growth in SCID (Severe Combined Immunodeficiency) mice injected with two different DKO clones (DKO-C1 and D1KO-C2) and EV cells. The upper panel was the representative tumor picture at the final day. The panel below was the tumor growth curve, which was shown as tumor volume with different post implantation days. N=5 for each group. **G&H**. RT-qPCR of *FGF18* expression and hemoglobin genes (*HBZ*, *HBE1* and *HBB*) expression upon siRNA knock-down in one of the DKO clone. Knocking down experiment was performed using two different siRNAs targeting *FGF18* gene and data was shown as the fold change relative to control siRNA, si-Scramble. **I**. Growth curve of normal K562 cells and siRNA knock-down in one of the DKO clone. Data was calculated as the fold change against day 0. All data shown here indicates average + standard error. P value was calculated by two-tailed student’s t-test. P value less than 0.05 was shown as *; P value less than 0.01 was shown as **; P value less than 0.001 was shown as ***.

First, we checked *FGF18* expression by RT-qPCR in different KO clones. S1KO and S2KO both showed *FGF18* expression upregulation compared to the empty vector (EV) (Figure 1B, Figure S1C), suggesting that both S1 and S2 can function as silencers although they may not have equivalent silencing capability because we observed they had different extents of gene upregulation levels upon silencer removal. Remarkably, DKO showed dramatic upregulation which was more than the sum of S1KO and S2KO (Figure 1B, Figure S1C) implicating the synergism between the two silencer components.

In our previous work, we showed that S1KO cells exhibited erythroid differentiation *in vitro* and tumor growth inhibition *in vivo* (Cai et al., 2021). Here, again, we observed cell adhesion of S2KO and DKO cells (Figure S1D), although S2KO and DKO showed similar cell adhesion ability to fibronectin as S1KO. We wanted to know whether S2KO and DKO cells exhibited similar phenotypes as S1KO cells and whether DKO confers synergistic effects on these phenotypes. To explore this, we overlapped the RNA-seq data of different knock-out clones and found that the triple intersected part of differentially expressed genes was significantly enriched with several cancer hallmarks including heme metabolism (Figure 1C) which is associated with erythroid differentiation phenotype that we previously elucidated in S1KO cells (Cai et al., 2021; Chiabrando, Vinchi, Fiorito, Mercurio, & Tolosano, 2014).

To further confirm this phenotype, we checked the erythroid differentiation indicators (hemoglobin genes: *HBZ*, *HBE1* and *HBB*) (Y. N. Ma et al., 2013) by RT-qPCR as we did previously (Cai et al., 2021). We found that DKO dramatically increased these erythroid differentiation markers compared to single knock-out clones (Figure 1D, Figure S1E), which is consistent with the synergistic upregulation of *FGF18* expression (Figure 1B, Figure S1C).

Leukemic erythroid differentiation can cause cell growth inhibition which is one of the existing therapies for leukemia (Hietakangas et al., 2003; Martin, Bradley, & Cotter, 1990). Therefore, we asked if DKO also shows synergistic effects in terms of growth inhibition. We performed the cell growth assay *in vitro* for different KO clones (Figure 1E, Figure S1F) and xenograft experiments *in vivo* by injecting two different DKO clones (clone1 of DKO, “DKO-C1” and clone2 of DKO, “DKO-C2”) into the mice (Figure 1F, xenograft experiments *in vivo* for two different S1KO clones see Figure D-E, Cai et al., 2021). Both pieces of data showed that DKO had synergistic growth inhibition *in vitro* and *in vivo*, which was in line with the synergistic erythroid differentiation phenotype of DKO.

To confirm that the phenotypes we observed were due to the upregulation of *FGF18* gene expression, we performed siRNA knock down experiments against the *FGF18* gene in DKO cells. We successfully knocked down the *FGF18* gene using two different siRNAs (Figure 1G) and by performing RT-qPCR against haemoglobin genes, we found that knocking down *FGF18* gene caused downregulation of haemoglobin genes indicating that the erythroid differentiation phenotype was partially because of the *FGF18* gene (Figure 1H). Moreover, we also performed the growth assay for the wild type K562 cells, DKO cells with siRNAs targeting *FGF18* and DKO cells with control siRNA (Figure 1I). Our results showed that knocking down *FGF18* in DKO cells can significantly increase the cell growth as compared with control DKO cells, although the cell growth rate was still lower than the wild type K562 cells (Figure 1I). These results showed that there is a direct relationship between *FGF18* gene and observed phenotypes, but other genes such as *SH3PXD2B* (Figure S1G) may also contribute to the phenotype since we observed that siRNAs against *FGF18* gene did not fully rescue back the CRISPR knockout phenotypes.

In summary, S1 and S2 can synergistically repress *FGF18* expression. DKO leads to both quantitative changes and qualitative changes including increased *FGF18* gene upregulation and more dramatic phenotypic changes such as erythroid differentiation and growth inhibition both *in vitro* and *in vivo*. These results indicate that S1 and S2 can function together to synergistically upregulate a target gene.

### 3D genome organization underlies the synergism of silencers

In the results above, we revealed an example of two distal silencers working together in synergy to boost target gene expression. Next, we asked if there is any interplay between S1 and S2, and whether there are any chromatin organization changes in KO cells. To investigated this, we first performed 4C-seq using *FGF18* promoter as the viewpoint in different clones (EV, S1KO and DKO) since chromatin interactions are the key determinants for modulating gene expression (Babu & Fullwood, 2015; Deng et al., 2012; Q. Li, Barkess, & Qian, 2006; Perino & Veenstra, 2016).

We found that silencers can stabilize the chromatin interaction landscape and their removal led to changes in the chromatin organization landscape (Figure 2A-B). Briefly, S1KO has 17 gained chromatin loops and 3 lost chromatin loops compared to control cells (Figure 2A). DKO lost 24 distant chromatin loops and gained 10 nearby chromatin loops compared to S1KO which further constrained all the chromatin loops to a narrow region (between *FGF18* gene and S2) to dramatically change the chromatin interaction landscape (Figure 2A-B). Consistent with our previous observations (Cai et al., 2021), S1KO and DKO both showed distal loops tended to change more easily than nearby loops (Figure S2A-B). Surprisingly, when we carefully dissected the loops in S1KO, we found there were multiple new loops formed around S2 site in S1KO cells (Figure 2B), suggesting that S2 can compensate the role of S1 through increased contact frequency to *FGF18* in S1KO. Overall, the data confirmed the interplay between different silencer components in terms of silencer functioning.

**Figure 2.**
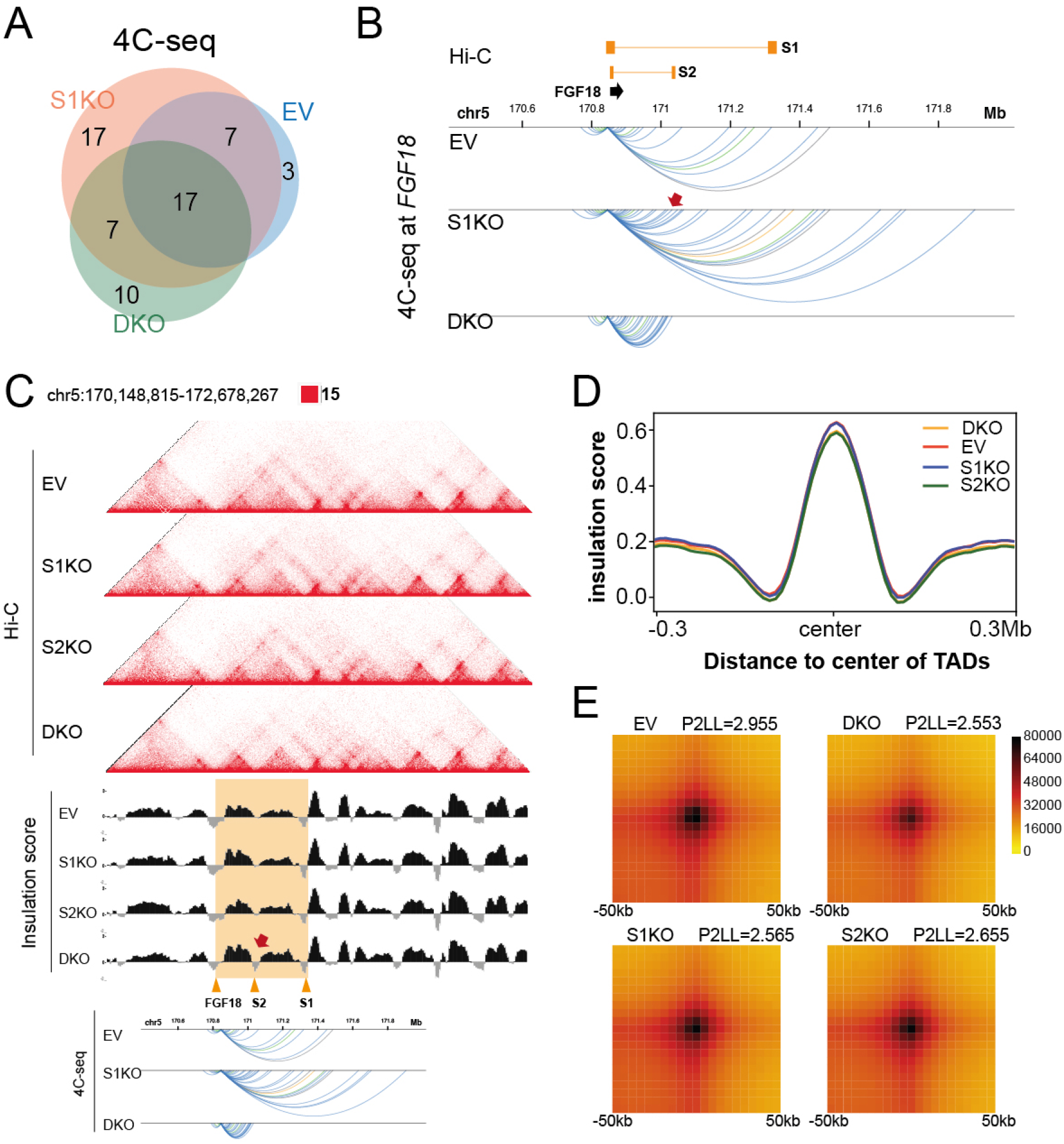
3D genome organization underlies the synergism of silencers. **A&B**. Venn diagram (A) and arcs (B) of significant chromatin interactions identified by 4C-seq using *FGF18* promoter as the viewpoint in EV, S1KO and DKO cells. Two replicates of 4C-seq were performed and significant chromatin interactions (p<0.05) analysed by R3Cseq package (Thongjuea, Stadhouders, Grosveld, Soler, & Lenhard, 2013) were shown in Venn diagram and arcs. *FGF18* gene region, S1 and S2 were indicated in **B**. The colors of the arcs represented the histone state of the chromatin interactions (yellow: H3K27ac associated loops; blue: H3K27me3 associated loops; green: both H3K27ac and H3K27me3 associated loops, grey: none of H3K27ac and H3K27me3 associated loops) (Cai et al., 2021). The new chromatin interaction to S2 site in S1KO was indicated by the red arrow. **C**. Screenshot showed the aligned Hi-C matrix of EV, S1KO, S2KO and DKO, insulation score of EV, S1KO, S2KO and DKO, and 4C-seq of EV, S1KO and DKO at the *FGF18* MRR region. *FGF18* MRR region was highlighted by orange box and some specific sites (FGF18 gene region, S2 and S1) were indicated as well. The new peak of insulation score at the S2 site in DKO cells was indicated by the red arrow. **D**. Mean plot described genome-wide insulation score around the TADs (use TADs in EV as the reference TADs) in EV and different KO cells (S1KO, S2KO and DKO). The X-axis represents the genome distance to the center of TADs, while the Y axis represented the insulation score. **E**. Aggregate peak analysis (APA) for all the loops in EV, S1KO, S2KO and DKO cells (use EV loops as the reference). Loops were aggregated at the center of a 50kb window in 5kb resolution. The ration of signal at the peak signal enrichment (P) to the average signal at the lower left corner of the plot (LL) (P2LL) were indicated to show the normalized intensity of all the loops.

To further characterize whether the TAD and loop changes occurred locally at the *FGF18* silencer region or genome-wide, we performed Hi-C in different KO cells (EV, S1KO, S2KO and DKO cells) and found that the number of TADs and loops were similar in different KO cells (Figure S2C), suggesting changes in the genome-wide TADs and loops changes were subtle.

In contrast, at the *FGF18* silencer region, we observed changes in the Hi-C matrices where the loop between *FGF18* gene promoter and S1 was lost and the sub-TAD between *FGF18* gene promoter and S2 was stronger (Figure 2C). To further confirm such changes, we calculated the TAD insulation scores for the different KO cells which stand for the strength of the TADs. Consistent with the Hi-C matrices, a new TAD boundary was identified at the S2 site in the DKO cells, which again indicated the formation of a stronger sub-TAD formation between *FGF18* gene promoter and S2 site (Figure 2C, Figure S2D). This was also concordant with the 4C-seq data which showed that all the chromatin interactions in the DKO cells were constrained inside of this strong sub-TAD region (Figure 2C).

Apart from the *FGF18* silencer region, other genomic regions did not show any obvious changes. To explore whether the TADs and loops changes at *FGF18* silencers region were local or in the genome-wide, we calculated the insulation score for all the TADs in different KO cells (Figure 2D). Data showed that the average insulation scores for all the TADs in different samples were highly similar (Figure 2D). We also analysed the loop strength by aggregate peak analysis (APA) plot for different KO cells, using the P2LL score (the ration of signal at the peak signal enrichment (P) to the average signal at the lower left corner of the plot (LL)) as a measure of loop strength (Figure 2E). The results showed that the loop strength was highly similar across all four samples (Figure 2E). The Hi-C matrix of another MRR region at the *IGF2* gene, which was previously confirmed to be a silencer region (Cai et al., 2021), did not show any obvious changes of Hi-C matrix (Figure S2E), suggesting that our KO of the silencers of *FGF18* did not affect the 3D genome organization of any other MRR regions. Together with the insulation score analysis for the TADs and APA plot, our results suggested that changes for the loops and TADs at the *FGF18* region were local and specific, while other genomic regions remained highly unchanged.

Taken together, we showed that the silencers S1 and S2 interact and work together to form a super-silencer and knocking out silencers will destabilize the genome organization by affecting the TADs and loops locally. These results suggest that at least one mechanism of action of super-silencers is that the silencers components interact with each other via chromatin interactions to regulate each other and their target genes.

### Epigenomic differences together with chromatin interactions underlie the action of “super-silencer”

Besides the 3D genome organization changes, we further explored the epigenomic differences between the S1KO and DKO to investigate the detailed synergistic mechanism because histone marks are another epigenetic aspect that can modulate gene expression (Bannister & Kouzarides, 2011; Karlic, Chung, Lasserre, Vlahovicek, & Vingron, 2010). We performed H3K27ac ChIP-seq and H3K27me3 ChIP-seq in different KO cells (EV, S1KO and DKO). Specifically at the *FGF18* gene region, there were more H3K27ac peaks and fewer H3K27me3 peaks in DKO compared with S1KO and EV, which could explain the greater upregulation of *FGF18* in DKO (Figure 3A). The increased H3K27ac signals were further confirmed by the ChIP-qPCR at five different regions (R1-R5) along the *FGF18* gene body (Figure 3A).

**Figure 3.**
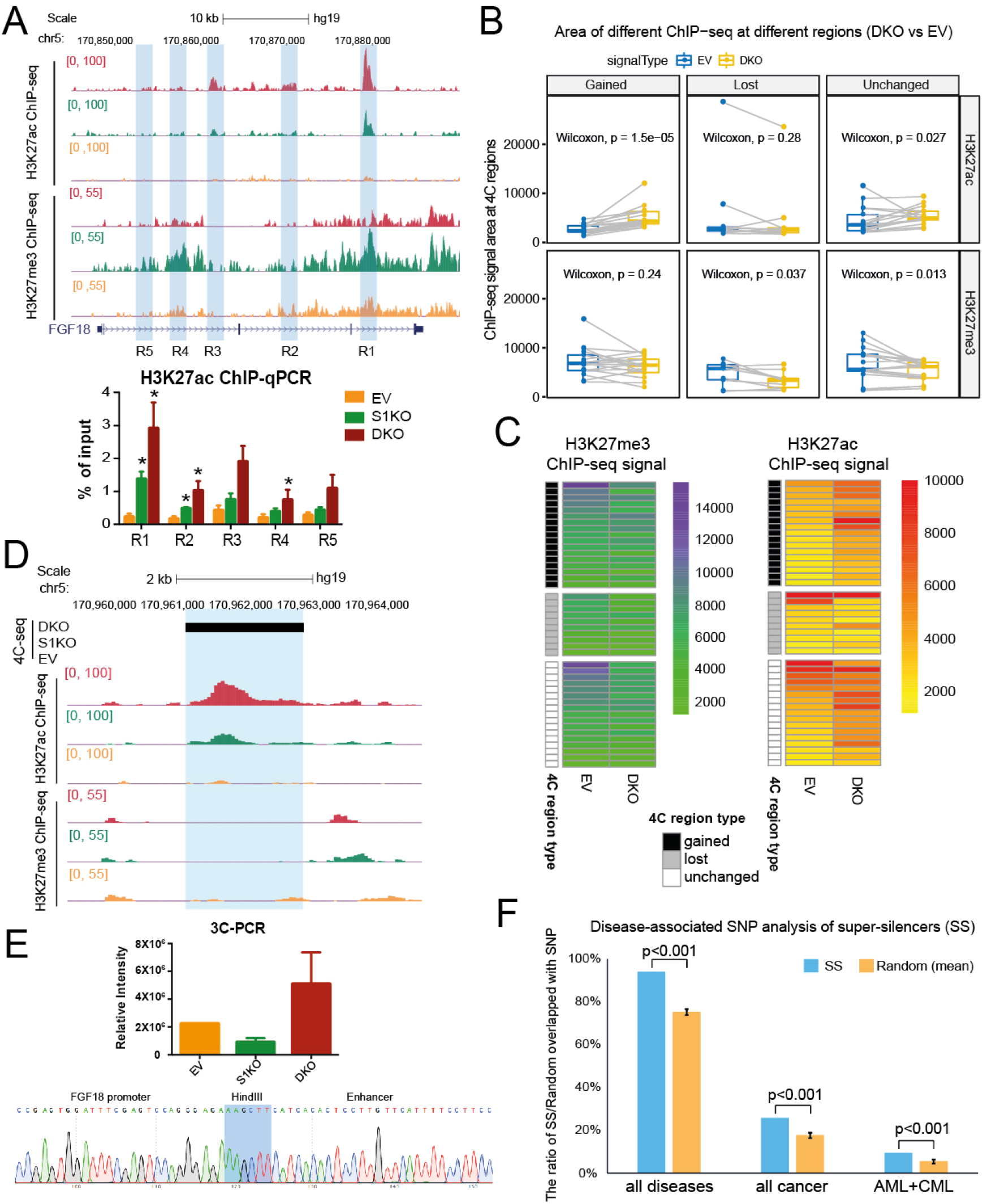
Unchanged loops and gained loops to *FGF18* in DKO became increased H3K27ac signals and decreased H3K27me3 signals. **A**. H3K27ac ChIP-seq and H3K27me3 ChIP-seq of EV, S1KO and DKO cells were shown at the FGF18 gene region. Five regions along the *FGF18* gene body (R1-R5) was performed ChIP-qPCR against H3K27ac marks, which were shown as % of input. ChIP-seq and ChIP-qPCR were performed by three different replicates. Data shown here were average + standard error. P value was calculated by two-tailed student’s t-test. P value less than 0.05 was shown as *. **B**. Boxplots of ChIP-seq signal changes of H3K27me3 and H3K27ac at different types of 4C regions (gained, lost and unchanged 4C loops) in EV and DKO cells. The same 4C regions are connected by gray lines. P values were indicated in each boxplot. **C**. Heatmap of ChIP-seq signal changes of H3K27me3 and H3K27ac at different types of 4C regions (gained, lost and unchanged 4C loops) in EV and DKO cells. **D**. Screenshot described one of the new enhancers in DKO cells showing H3K27ac ChIP-seq, H3K27me3 ChIP-seq and 4C-seq in EV, S1KO and DKO cells. The new enhancer was highlighted by blue. **E**. 3C-PCR of the new enhancer described in **D** of EV, S1KO and DKO cells by two independent 3C libraries. Data shown here was relative intensity measure by ImageJ. The panel below was the sanger sequencing results for the 3C ligated fragment. The *FGF18* promoter sequence, HindIII cut site and the new enhancer sequence were indicated. **F**. Disease-associated Single Nucleotide Polymorphisms (SNPs) analysis of different groups (all diseases, all cancer and AML+CML) with super-silencers (SS). SNPs was overlapped either with super-silencers or the random regions, and the y axis showed the ratio of SS/Random overlapped with different SNP groups. P values were indicated for each group comparing super-silencers with random regions.

To systematically and deeply dissect the synergistic mechanism in depth, we performed the integrative analysis of 4C-seq and ChIP-seq. We classified the loops into three different categories including gained loops, lost loops and unchanged loops and correlated them with histone modifications. We found that there were increased H3K27ac signals and decreased H3K27me3 signals for unchanged loops in DKO compared with EV (Figure 3B-C). Moreover, H3K27ac signals were also increased at gained loops (Figure 3B-C), suggesting that loops were more active in DKO cells. Gained H3K27ac signals and depleted of H3K27me3 signals again was observed when comparing DKO vs S1KO (Figure S3A-B) but not observed when comparing S1KO vs EV (Figure S3C-D). This suggested that there are indeed differences between S1KO to DKO. As for S1KO, there were not many epigenomic differences observed, possibly due to the compensation by S2.

The increased H3K27ac signals in DKO gained loops (Figure 3B, Figure S3A) suggested that there might be new enhancers which looped to *FGF18* gene promoter in DKO. Indeed, we observed multiple distal new enhancers that can activate *FGF18* expression in DKO (Figure 3D, Figure S3E). One new enhancer which looped to *FGF18* gene promoter only in DKO was validated by 3C-PCR and Sanger Sequencing, confirming increased contact frequency between the new enhancer and the *FGF18* gene promoter in DKO (Figure 3D-E).

Our previous work concluded that initial histone states can predict the changed loops upon *IGF2* looping silencer removal (Cai et al., 2021). Here we performed a similar analysis. However, for different initial histone states (high, medium and low), we did not observe any consistent trends in loop changes (Figure S3F). Just as different SEs work via different mechanisms, the *FGF18* silencers may work through a different mechanisms as compared to the *IGF2* looping silencer. It will be interesting to explore whether the conclusion that initial histone states can predict the changes of chromatin interactions is common in different cellular backgrounds. Taken together, our results showed that changes in histone modifications also occur upon silencer knockout, and these may work together with changes in chromatin interactions. We suggest that the combined changes of histone modifications and chromatin interactions may lead to the synergistic upregulation of *FGF18* in DKO.

This work is, to the best of our knowledge, the first indication that two distal silencers can cooperate to function as “super-silencers”. Next, we asked what is the functional importance of MRRs, our putative “super-silencers”, in terms of their relationship to disease. Disease-associated sequence variation has been shown to be enriched in SEs (Hnisz et al., 2013). Therefore, we performed a similar analysis to explore the relationship between super-silencers and single nucleotide polymorphisms (SNPs) in different disease groups (all diseases, all cancer and Acute myeloid leukemia (AML)+Chronic myeloid leukemia (CML)). Results showed that compared to the random regions, super-silencers (SS) were enriched for SNPs associated with all diseases, all cancer and AML+CML groups (Figure 3F). The ratio of SNPs in different groups (all diseases, all cancer and AML+CML) overlapped with SS showed the similar results (Figure S3G). This analysis highlighted the importance of super-silencers in terms of diseases such as cancers, and elucidated the significance of targeting super-silencers to treat cancers.

### Combinational treatment of GSK343 and X5050 leads to synergistic loss of TAD and loops

MRRs are enriched with H3K27me3 signals, therefore, we asked whether the removal of H3K27me3 signals will affect the 3D genome architecture of super-silencers. We applied a methyltransferase inhibitor, GSK343 (5uM), to deplete H3K27me3 histone modifications. Previously, we performed 4C-seq upon GSK343 treatment and found that only long-range chromatin interactions altered, while short-range chromatin interactions remained unchanged (Cai et al., 2021). To characterize the genome-wide TADs and loops, here we performed Hi-C analysis and found modest changes in overall TADs and loops upon GSK343 treatment (Figure 4A-B, Figure S4A, Figure S4C).

**Figure 4.**
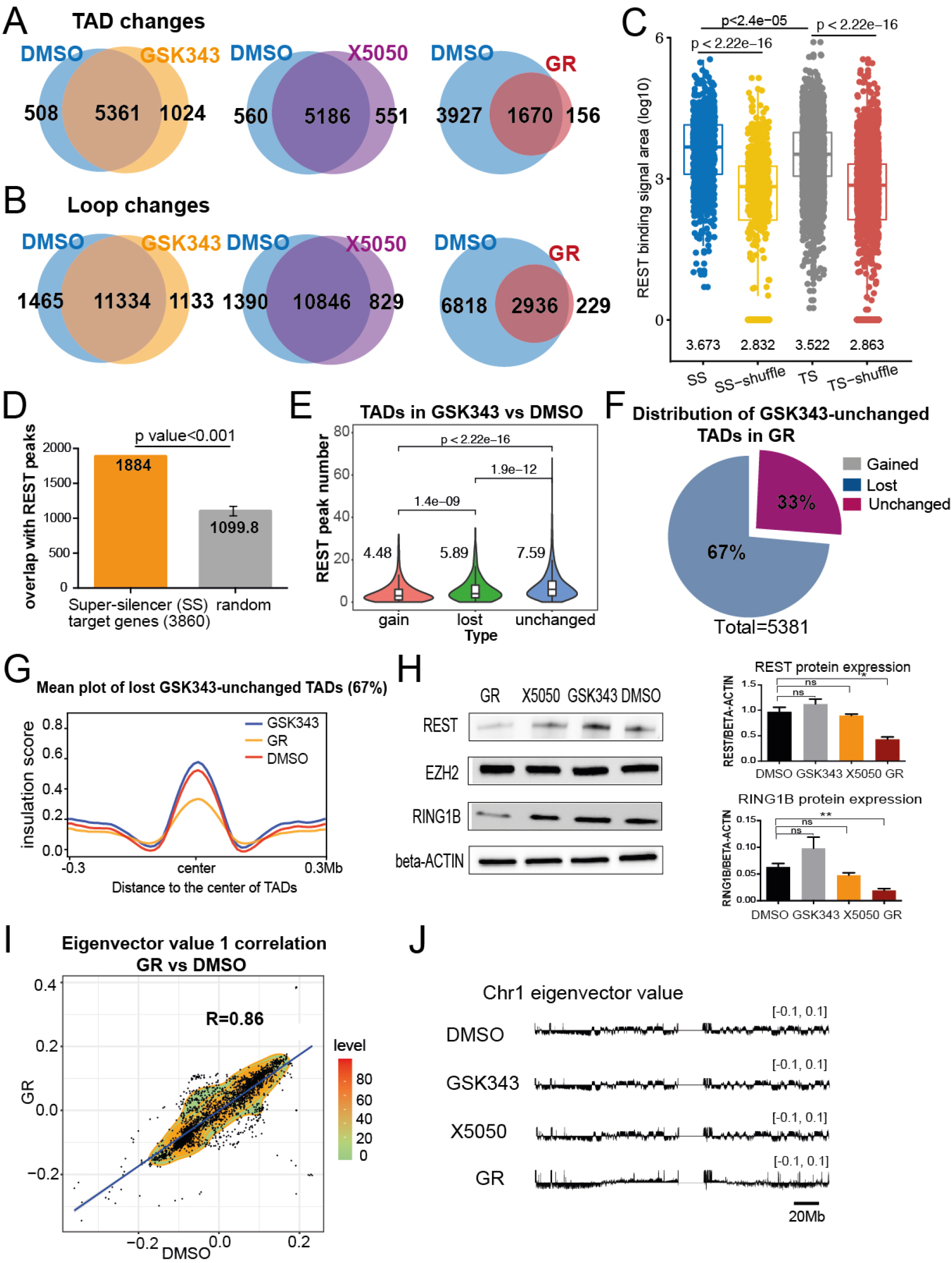
Combinational treatment of GSK343 and X5050 led to synergistic loss of TAD and loops. **A**. Venn diagram of TAD changes in DMSO, GSK343, X5050 and combinational treatment of GSK343 and X5050 (“GR”). **B**. Venn diagram of loop changes in DMSO, GSK343, X5050 and combinational treatment of GSK343 and X5050 (“GR”). **C**. Boxplot described the REST binding enrichment at super-silencers (SS) and typical silencers (TS). SS-shuffle and TS-shuffle serve as the random control. REST enrichment was shown as REST binding signal area (log10). **D**. REST enrichment at the super-silencer (SS) target genes and random genomic regions. The y axis showed that overlapped number between the SS target genes/random and REST ChIP-seq peaks. **E**. Plot described the REST peak numbers at different TAD categories (gained, lost and unchanged) in GSK343 vs DMSO. **F**. Distribution of the unchanged TADs upon GSK343 in the combinational treatment of GSK343 and X5050 (“GR”). **G**. Mean plot described genome-wide insulation score around the TADs of 67% lost GSK343-unchanged TADs in the combinational treatment (“GR”). The X-axis represents the genome distance to the center of TADs, while the Y axis represented the insulation score. **H**. Western blot showed protein expressions of REST, EZH2, RING1B and beta-actin in DMSO, GSK343-only, X5050-only and GR conditions. The protein level measurement of REST and RING1B were performed by ImageJ against beta-actin levels in different conditions. NS stands for no significance; P value less than 0.05 was shown as *; P value less than 0.01 was shown as **. **I**. Density plot described the global correlation between the eigenvector value from DMSO condition and combinational treatment (“GR”) condition at 1Mb resolution. The X axis represents the eigenvector value in the DMSO condition, while the Y axis represents the value in the GR condition in the same locus. **J**. Representative eigenvector value for 50kb resolution at chromosome 1 in DMSO, GSK343-only, X5050-only and GR conditions.

We reasoned that just as SEs are associated with multiple transcription factors, super-silencers might also be associated with multiple transcription factors and multiple epigenetic drugs targeting different transcription factors would need to be targeted in order to change the super-silencers and chromatin interactions. As a first step in identifying additional factors to target, we wished to elucidate the super-silencer-controlled genes. To find the potential super-silencer-controlled genes such as *FGF18* gene, we performed H3K27me3 HiChIP experiment in normal K562 cells to pulldown the H3K27me3 marks and identify the genome-wide chromatin interactions at the same time. We identified 3860 genes associated with two or more H3K27me3 HiChIP loops, similar to *FGF18* gene, and termed these genes as potential “super-silencer target genes”.

Next, we examined super-silencers and super-silencer target genes for associated transcription factors. We found that REST is enriched in super-silencers compared with random regions and typical silencers (Figure 4C). REST is known to be a repressor to repress neuron gene expression (Hwang & Zukin, 2018). Furthermore, we found that REST was enriched at these potential super-silencer target genes (Figure 4D) when compared to the random regions. Moreover, REST was more enriched at TADs that remained unchanged after GSK343 treatment, compared to gained and lost TADs (Figure 4E), suggesting that REST may be the factor that could help to retain the structure TADs upon GSK343 treatment. We reasoned that applying REST inhibitor together with GSK343 may disrupt the structures of TADs and loops.

As a first step, we asked whether REST is directly involved in the TADs formation and super-silencer functioning. We applied a REST inhibitor, X5050 (Charbord et al., 2013), to K562 cells and performed Hi-C. However, similar to the GSK343 treatment, the overall TADs and loops did not change a lot (Figure 4A-B, Figure S4B, Figure S4D), suggesting that similar to H3K27me3 marks, targeting REST alone cannot ablate TAD structures.

In contrast, Hi-C analysis showed that the combinational treatment of GSK343 and X5050 together led to severe loss of TADs and loops (Figure 4A-B, Figure S4E-F). Specifically, among TADs that were unchanged after single GSK343 treatment, GR treatment led to the loss of 67% of these TADs (Figure 4F) together with decreased TAD insulation score (Figure 4G).

Interestingly, upon performing western blot of REST protein for the conditions of REST inhibition alone and GR, we found that X5050 does not serve as a REST degrader by its own in K562 cells (Figure 4H). This observation was confirmed using two different REST antibodies (Figure 4H, Figure S4H). Instead, it functions as a REST degrader only in the GSK343 sensitized environment (Figure 4H), which might be one of the reasons why GR can cause severe changes of genome architecture, while REST inhibition alone in K562 led to fewer changes.

We also tested the protein expression of PRC2 subunit EZH2 (van Mierlo, Veenstra, Vermeulen, & Marks, 2019) and PRC1 subunit RING1B (Stock et al., 2007), and found that RING1B expression was decreased in the GR condition while EZH2 remained unchanged for all the conditions (Figure 4H, Figure S4I). Given that similar to REST, RING1B was also enriched at the unchanged TADs upon GSK343 (Figure S4G), RING1B may play role for the GR synergism as well. Besides the loss of TADs and loops, GR also showed altered A/B compartments (Figure 4I-J) while GSK343 and X5050 treatment alone did not shown obvious changes in A/B compartments (Figure S4J-K), suggesting that there are alterations in transcription in the GR condition.

### The combinational treatment of GSK343 and X5050 shows synergistic antitumor effects

Since combinational treatment of GSK343 and X5050 (GR) led to severe loss of TADs and loops and alterations of A/B compartments, we hypothesized that GR will give rise to transcriptional dysregulation, which may affect the cell identity. To test this, first, we performed the dose response matrix of different concentrations of GSK343 and X5050, and calculated the Bliss synergy score to be 17.899 (Figure 5A). The Bliss synergy score corresponds to the excess response due to drug interactions, and a score larger than 10 represents high likelihood for synergism.

**Figure 5.**
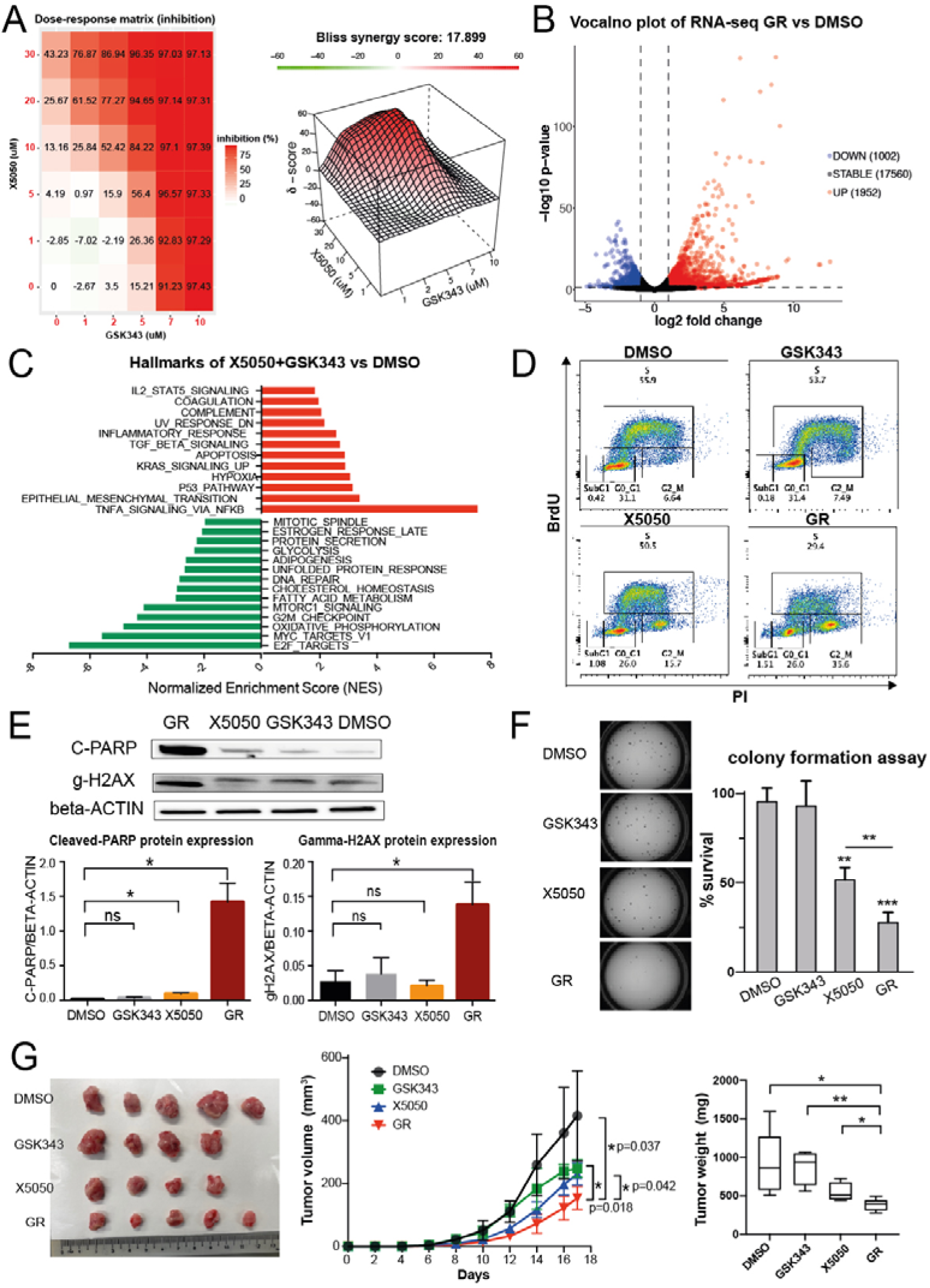
Combinational treatment of GSK343 and X5050 had synergistic antitumor effects. **A**. Test the synergy of drug combinations using Bliss score. Different concentrations of GSK343 and X5050 in K562 cells were tested and the dose-response matrix was shown by % inhibition. Bliss score was calculated and the Bliss score larger than 10 indicated that two drugs were likely to be synergistic. **B**. Vocalno plot of RNA-seq comparing GR versus DMSO condition. The X axis showed the log2 fold change, while the Y axis showed the −log10 p-value. The number of downregulated, stable and upregulated genes were indicated. **C**. Gene Set Enrichment Analysis (GSEA) was performed using the differentially expressed genes in GR compared with DMSO against cancer hallmark database (Subramanian et al., 2005). Significant upregulated and downregulated terms were shown here and ranked by Normalized Enrichment Score (NES). **D**. Cell cycle analysis was performed in different drug treated conditions (DMSO, GSK343-only, X5050-only and GR) by BrdU labelling. The X axis showed the signal of propidium iodide (PI), while the Y axis showed the signal of BrdU in the flow cytometry analysis. The percentage of subG1, G0/G1, S and G2/M phases were indicated for different drug treated conditions. **E**. Western blot showed protein expressions of cleaved-PARP (C-PARP), gamma-H2AX (g-H2AX), and beta-actin in DMSO, GSK343-only, X5050-only and GR conditions. The protein level measurement of cleaved-PARP and gamma-H2AX were performed by ImageJ against beta-actin levels in different conditions. **F**. Colony formation assay of K562 cells in DMSO, GSK343-only, X5050-only and GR conditions. Representative pictures and percentage of survival were shown. **G**. Tumor growth in NSG mice (NOD scid gamma mice) injected with K562 cells together with different drugs (DMSO, GSK343-only, X5050-only and GR). The left panel was the representative tumor picture at the final day. The panel in the middle was the tumor growth curve, which was shown as tumor volume (mm^3^) with different post implantation days. The right panel was the tumor weight (mg) at the final day. N=5 for each group. All data shown here indicates average +/− standard error. P value less than 0.05 was shown as *, P value less than 0.01 was shown as **.

To explore the transcriptional changes in single and combinational treatment, we performed RNA-seq for the DMSO, GSK343-only, X5050-only and GR conditions. The results showed that X5050-only generated mild changes of gene expressions (Figure S5B), while GSK343-only can cause a lot of gene expression changes (Figure S5A) and GR had the most dramatic changes of gene expression (Figure 5B). To explore the phenotype after drug treatment, we performed the gene set enrichment analysis (GSEA) for cancer hallmarks in the GR condition (Figure 5C). Terms such as “Apoptosis” and “P53 pathway” were significantly enriched in the GR condition, while terms such as “G2M checkpoint” and “DNA repair” were depleted in the GR condition (Figure 5C). To further test these processes, we performed cell cycle analysis through flow cytometry, and found that X5050-only showed a mild decrease in the number of cells in S phase and mild increase in the number of cells in G2M phase, while GR showed an obvious decrease in the number of cells in S phase and increase in the number of cells in G2M phase (Figure 5D). This result was consistent with the downregulation observed at G2M checkpoint related genes (Figure 5C, Figure S5C).

To test whether the GR treatment led to apoptosis and DNA damage, which were showed by the hallmark analysis (Figure 5C, Figure S5D-E), we performed western blot for the DMSO, GSK343-only, X5050-only and GR conditions. Cleaved-PARP, which is a marker for the apoptosis (Kaufmann, Desnoyers, Ottaviano, Davidson, & Poirier, 1993), showed a mild increase in the X5050-only condition and a significant, dramatic increase in the GR condition, suggesting more apoptosis was happening the GR condition (Figure 5E). Gamma-H2AX, which is a marker for the DNA damage signals (Mah, El-Osta, & Karagiannis, 2010), showed no significant difference in the GSK343-only condition and X5050-only condition but showed a significant increase in the GR condition, suggesting that there is significant DNA damage occurring in the GR condition (Figure 5E).

The results above indicated that GR condition had synergistic effects in terms of G2M cell cycle changes, increased apoptosis and increased DNA damage, which are all processes that lead to reduction in tumor progression. Therefore, we wondered whether GR could have antitumor effects. To test this, first, we performed the colony formation assay in K562 cells for DMSO, GSK343-only, X5050-only and GR conditions (Figure 5F). Data showed that GSK343 itself did not cause any changes of colony formation, while X5050 single treatment can reduce some colony formation ability (Figure 5F). GR exhibited more dramatic colony formation inhibition as compared with the X5050 single treatment, suggesting GR can synergistically reduce colony formation ability. Next, we performed the *in vivo* experiments by inoculating K562 cells into the mice and injecting different drugs (DMSO, GSK343-only, X5050-only and GR) (Figure 5G). Consistent with the colony formation assay, GSK343-only did not show any tumor volume reduction, while X5050-only reduced the tumor volume to some extent (Figure 5G). GR showed the most dramatic tumor growth inhibition as compared to other conditions, indicating the synergistic antitumor effects of GR (Figure 5G).

Together, combinational treatment of GSK343 and X5050 can work synergistically including cell growth inhibition, cell cycle arrest, colony formation inhibition and tumor growth inhibition, suggesting the potential usage of GR in treating cancer.

### Super-silencers are susceptible to the combinational treatment of GSK343 and X5050, which leads to upregulation of super-silencer-controlled apoptotic genes

We observed that combinational treatment of GSK343 and X5050 led to 67% loss of TADs, thus, we asked whether these lost TADs are associated with super-silencers. To test this question, we associated the H3K27me3 marks, super-silencers and H3K27ac marks with the lost TADs. Upon GSK343-only treatment, more H3K27ac-related-TADs lost compared to H3K27me3-related-TADs and super-silencers-related-TADs (Figure 6A). Upon X5050-only treatment and GR treatment, more H3K27me3-related-TADs and SS-related-TADs were lost compared to H3K27ac-related-TADs, suggesting that these TADs with H3K27me3 marks and super-silencers were more susceptible to be lost upon GR (Figure 6A). Previously, we showed that GR displayed more apoptosis (Figure 5E). We also found that SS controlled genes are largely overlapped with the apoptosis related genes (Figure 6B). Therefore, we hypothesized that GR treatment disrupted the super-silencers, which led to upregulation of targeted apoptosis genes and gave rise to the apoptosis phenotype.

**Figure 6.**
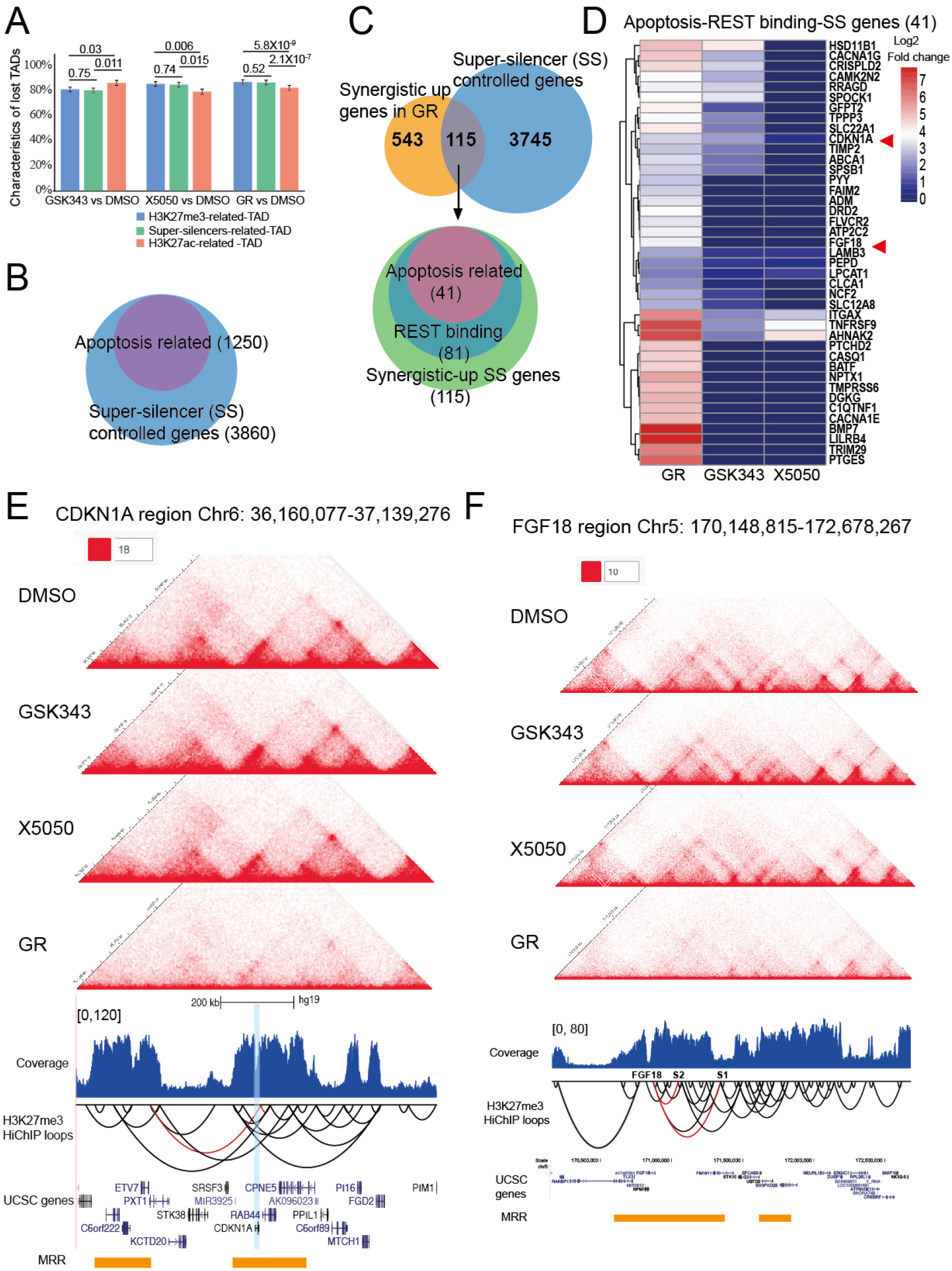
Combinational treatment of GSK343 and X5050 led to upregulation of super-silencers-controlled apoptotic genes. **A**. Characteristics of lost TADs in different conditions (GSK343 vs DMSO, X5050 vs DMSO, and GR vs DMSO). Three different groups of TADs were characterized including H3K27me3-related-TADs (TADs overlapped with H3K27me3 marks), super-silencers-related-TADs (TADs overlapped with super-silencers) and H3K27ac-related-TADs (TADs overlapped with H3K27ac). P values were indicated for the comparison of different groups. **B**. The overlap of apoptosis related genes with the super-silencer (SS) controlled genes. **C**. Schematic described the large overlap of the REST binding apoptosis associated genes with the synergistic upregulation SS-controlled genes. Numbers in each category were indicated. **D**. Heatmap showed the gene expressions of 41 shortlisted apoptosis-REST binding-SS genes in **C**. Gene expression were shown by log2 fold change in different conditions (GR, GSK343-only and X5050-only) against DMSO condition. Two interesting genes were indicated by the red arrow, which had screenshots in **E** and **F**. **E**. Screenshot at the *CDKN1A* region showed the Hi-C data in different conditions (DMSO, GSK343-only, X5050-only and GR), H3K27me3 HiChIP data including coverage and loops in normal K562 cells, UCSC gene tracks and MRR annotations. *CDKN1A* gene was highlighted and the H3K27me3 HiChIP loops associated with *CDKN1A* gene were indicated by red color. **F**. Screenshot at the *FGF18* region showed the Hi-C data in different conditions (DMSO, GSK343-only, X5050-only and GR), H3K27me3 HiChIP data including coverage and loops in normal K562 cells, UCSC gene tracks and MRR annotations. The location of *FGF18* gene, S1 and S2 were indicated, and the H3K27me3 HiChIP loops associated with *FGF18* gene, S1 and S2 were indicated by red color.

To find the potential GR controlled and SS controlled apoptosis genes, we first shortlisted the synergistically upregulated genes in GR, which were upregulated in excess to the sum of upregulation by single drugs. Next, we overlapped these 658 genes with the potentially SS controlled genes identified by the H3K27me3 HiChIP, and found 115 synergistic-up SS genes (Figure 6C). Among these genes, 81 genes were associated with REST binding and 41 genes out of these 81 genes were associated with apoptosis (Figure 6C), suggesting that synergistic upregulation genes upon GR were largely bound by REST and they largely control apoptotic processes. The final apoptosis-related REST binding super-silencer-associated synergistic upregulated genes were shown in Figure 6D by log2 fold change in GR, GSK343-only and X5050-only conditions. Among these 41 apoptosis-REST binding-SS genes, two potential interesting genes were indicated by red arrows: *CDKN1A* gene and *FGF18* gene (Figure 6D), whose synergistic upregulation in GR were confirmed by RT-qPCR (Figure S6A-B).

Cyclin-dependent kinase inhibitor 1 (*CDKN1A*), also known as p21, is one of the major targets of p53 that mediates cell cycle arrest and apoptosis (Karimian, Ahmadi, & Yousefi, 2016). Here we found that *CDKN1A* gene belongs to one MRR in K562 cells and regulated by two H3K27me3 HiChIP loops connecting to two different MRRs, suggesting that *CDKN1A* gene is regulated by “super-silencers” (Figure 6E). Upon GR treatment, chromatin interactions that connect the “super-silencers” with target genes are disrupted as shown by the Hi-C, leading to increased gene transcription (Figure 6D, Figure S6A). *CDKN1A* gene could be one of the genes in this category. Our results show that upon GR treatment, the super-silencers were disrupted while the targeted genes were upregulated, and these include cell cycle and apoptosis-related genes, which could explain the cell cycle arrest and apoptosis phenotypes observed upon GR treatment.

Another example is the *FGF18* gene, which we previously showed to be controlled by a super-silencer (Figure 6F). Previously, we showed that upon double knock-out of two silencer components, *FGF18* gene was synergistically upregulated, with local changes of TADs showing the gain of sub-TADs (Figure 2C). Here we confirmed that the two silencer components show chromatin interactions to each other and *FGF18* gene by H3K27me3 HiChIP (Figure 6F). Upon GR treatment, the TADs and loops at this *FGF18* super-silencer region were largely lost (Figure 6F), which again supports our conclusion that upon GR treatment, the super-silencers were disrupted while the targeted genes were upregulated, which gave rise to cell cycle arrest and apoptosis phenotypes since *FGF18* has been shown to promote apoptosis (Portela et al., 2015).

Taken together, through the genome-wide intersection and two examples above, we demonstrated that GR treatment mediated apoptosis and cell cycle arrest may through disrupting the super-silencers and their associated chromatin interactions. Interestingly, GR treatment does not only affect chromatin interactions at super-silencers, but also chromatin interactions at other regulatory elements such as H3K27ac associated elements, although this is to a smaller extent than super-silencers (Figure 6A). Because H3K27ac and H3K27me3 are marks that can occur at the same amino acid of histone 3 and show interplay with each other (Zhang, Cooper, & Brockdorff, 2015), we reason that these changes may be due to interplay between super-silencers and other regulatory regions.

### Loss of TADs upon GR is not dependent on transcription but decreased CTCF protein

Next, we asked why GR treatment can cause such a dramatic loss of TADs and chromatin interactions. Recently, a drug called curaxin was reported to kill cancer cells and disrupt TADs (Kantidze et al., 2019). Studies about curaxins showed that the changes of TADs causing by curaxin were not dependent on the transcription, nor reduced CTCF protein levels, but rather due to the weaker binding of CTCF (Kantidze et al., 2019). Therefore, first, we wanted to explore whether changes in transcription resulted in the loss of TADs in the GR condition.

To do this, we selected the gene desert regions from the whole genome and demonstrated that the gene desert regions indeed showed very low transcription levels (Figure 7A). Next, by assigning the TADs into either the “all genes” category or the “gene deserts” category and calculating the insulation score changes in GR vs DMSO for all the TADs and TADs in the gene deserts (Figure 7B), we found that there were no significant differences between these two categories. This result indicated that even in the gene deserts there was also loss of TADs. One representative gene desert in chromosome 1 is shown in Figure 7C, and the Hi-C results indicated that regardless of whether genomic regions contained gene deserts or active transcription regions, there was loss of TADs and chromatin interactions. Therefore, similar to the curaxin treatment, the loss of TADs caused by GR were not due to changes in transcription.

**Figure 7.**
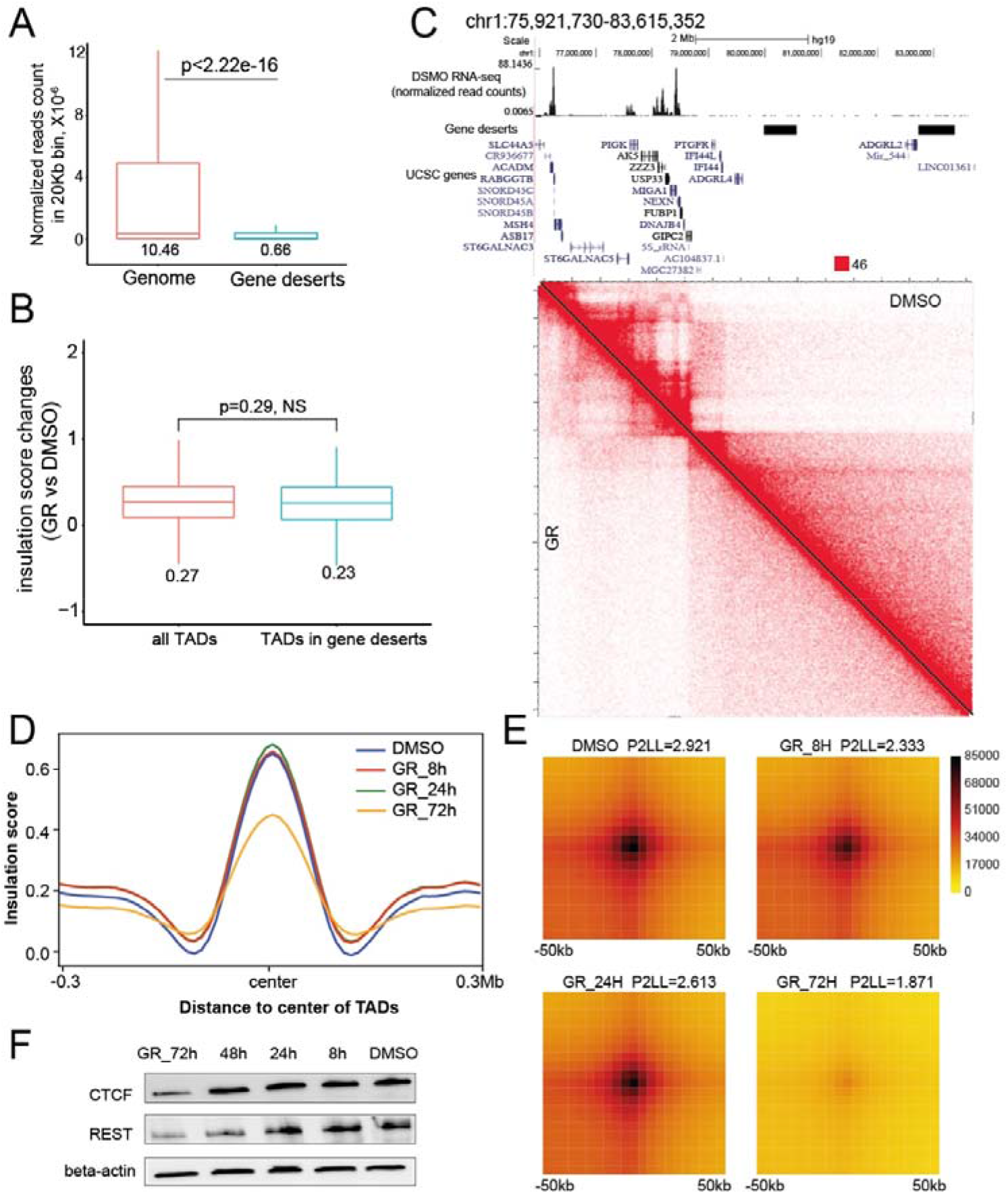
Loss of TADs upon GR do not dependent on transcription but decreased CTCF levels. **A**. Boxplot described the RNA normalized read count in a 20kb-long bin calculated from DMSO RNA-seq. Two categories of normalized read count were shown: genome and gene deserts, and p value was indicated for these two categories. **B**. Boxplot described the insulation score changes (GR vs DMSO) for either all TADs or TADs located in the gene deserts. P value was indicated for the comparison and NS stands for no significance. **C**. Genomic regions from chromosome 1 containing two representative gene deserts were shown. Screenshot contained DMSO RNA-seq shown as normalized read counts, gene deserts annotations, UCSC genes and corresponding Hi-C matrix in the DMSO and GR conditions. **D**. Mean plot described genome-wide insulation score around the TADs (use TADs in DMSO as the reference TADs) in DMSO and time course GR treated cells (GR_8h, GR_24h and GR_72h). The X-axis represents the genome distance to the center of TADs, while the Y axis represented the insulation score. **E**. Aggregate peak analysis (APA) for all the loops in DMSO and time course GR treated cells (GR_8h, GR_24h and GR_72h) (use DMSO loops as the reference). Loops were aggregated at the center of a 50kb window in 5kb resolution. The ration of signal at the peak signal enrichment (P) to the average signal at the lower left corner of the plot (LL) (P2LL) were indicated to show the normalized intensity of all the loops. **F**. Western blot showed protein expressions of CTCF, REST, and beta-actin in DMSO and time course GR treated cells (GR_8h, GR_24h and GR_72h).

The previous GR treatment was performed after 72 hours treatment, which was long enough to cause any secondary effects. To investigate changes resulting from GR treatment at earlier timepoints, we performed the time course treatment of GR in K562 cells including 8h, 24h and 72h. Then we performed Hi-C for different time points (Figure S7A), which showed that loss of TADs and loops only happened in the 72h treatment condition but not 8h and 24h conditions. We analysed the genome-wide TADs and loops for different condition, which again showed that decreased of insulation scores and loss of loops only happened in the 72h condition (Figure 7D-E). Two specific regions (*FGF18* super-silencer region and *CDKN1A* region) also supported this conclusion (Figure S7B-C). These results suggest that the changes in chromatin interaction were not a direct consequence of the drug changes but most likely due to secondary effects, unlike curaxins which can lead to changes in 3D genome organization within a few hours.

To explore the transcriptional changes, we performed the RNA-seq for the 8h and 24h conditions as well (Figure S7D-E), which exhibited similar numbers of differential expressed genes as the 72h (Figure 5B), suggesting that as early as 8h, the transcription levels already started to change. Again, the RNA-seq data supported the conclusion that although transcription levels has changed, the 3D genome organization still remained highly similar.

Next, we asked what secondary effects might lead to changes in 3D genome organization. Architecture proteins such as CTCF, RAD21 and SMC1A play important roles to maintain the TADs and loops (Narendra, Bulajic, Dekker, Mazzoni, & Reinberg, 2016; Nora et al., 2017; Wutz et al., 2017). Therefore, we checked the mRNA expression of these proteins by RT-qPCR, which showed that RAD21 and SMC1A levels remained unchanged while CTCF mRNA started to decrease after 24h treatment, but only exhibited significant decrease after 72h (Figure S7F). We further validated CTCF protein levels by western blot, which agreed with the RT-qPCR showing that the decrease of CTCF protein levels only started to be observed in 72h treatment (Figure 7F). Moreover, REST protein only decreased in the 48h and 72h conditions, while in 8h and 24 conditions, it still remained unchanged (Figure 7F). The decreased REST protein in the 72h condition might be the reason for the decreased CTCF levels and reduced CTCF levels of 72h condition might explain the loss of TADs and loops in the Hi-C experiment.

## Discussion

In this paper, we demonstrated the first results showing synergism between different silencers and that they can cooperate to form “super-silencers”. The debate as to whether super-enhancers are distinct entities from enhancers has not been settled yet and it is unclear whether constituent enhancers of super-enhancers cooperate synergistically or work individually (Frankel et al., 2010; Pennacchio et al., 2013). Similar questions could be raised with silencers. The synergistic collaborations between *FGF18* silencers would suggest that at least some silencers can act as “super-silencers”. Here we define “super-silencer” as a genomic region of high H3K27me3 signal comprising multiple silencers that can work as a whole entity to repress transcription of genes involved in cell identity. However, we cannot rule out the possibility that different “super-silencers” can work in different modes in other regions or cell lines, or maybe the working mode could be context dependent. Therefore, additional dissection about silencers is needed in the future to address the difference between silencers and “super-silencers”.

The relationship between 3D genome organization and transcription has been discussed for a long time (van Steensel & Furlong, 2019). 3D genome organization such as TADs are thought to regulate transcription: TAD boundaries can function to insulate the communication between promoters and enhancers, and loss of TAD boundaries can thereby lead to dysregulation of genes (Flavahan et al., 2016; Lupianez et al., 2015). Transcription sometimes can in turn affect the 3D genome organization as suggested by the transcription perturbation study via triptolide to target inhibition and flavopiridol to block to RNA polymerase II elongation. Li et al found that upon transcription perturbation, the TAD border strength was reduced, while the inter-TAD interactions increased (L. Li et al., 2015). Another study in *Drosophila* by applying triptolide showed the similar conclusion and more dramatic changes in TADs (Rowley et al., 2017). Here, we showed that CRISPR knock-out of silencers only can change the local TADs but the overall TADs remained similar, although the transcription patterns altered in the KO cells. During the GR treatment experiments, we demonstrated that loss of TADs was not dependent on transcription. Moreover, the early time course GR treatment (8h and 24h) results showed that transcription levels changed early on, but this led to subtle changes of TADs and loops. Our results added on to the evidence that although transcription can affect the 3D genome organization in some contexts, in other contexts, alterations of 3D genome organization are not dependent on the transcription.

3D genome organization can be affected by various determinants including architectural proteins such as CTCF and cohesin (Narendra et al., 2016; Nora et al., 2017; Wutz et al., 2017) alteration of transcription factors (Kim & Shendure, 2019) and histone modifications (J. Huang, Marco, Pinello, & Yuan, 2015). We elucidated that upon GR treatment, the loss of TADs and loops may due to the decreased levels of CTCF proteins since CTCF is the major factor that defines the TAD boundary and its decrease can lead to weaker TAD boundary strength (Nora et al., 2017). However, in the GR conditions, we also observed obvious apoptosis phenotypes, alterations of transcription factors and changes in histone modifications, which may account for the loss of TADs and loops. It would be interesting to explore the relationship between the above changes and loss of TADs.

Recently, phase separation model has been proposed to link between the super-enhancers and the gene activation. Transcriptional coactivators such as BRD4, MED1 and OCT4 showed liquid-like condensates at the super-enhancers, suggesting a model that phase-separated condensates at the super-enhancers can concentrate the transcription apparatus to activate cell identity genes (Boija et al., 2018; Sabari et al., 2018). Besides activation, the heterochromatin protein 1 (HP1) which can mediate gene silencing also showed phase-separated droplets (Larson et al., 2017), indicating the phase separation may be a general mechanism which not only happens in the activation but also silencing process. Therefore, phase separation may one of the mechanisms underlying the functioning of human super-silencers, which needs to be further explored.

In conclusion, our data demonstrated the first example of a “super-silencer”, constitute of two silencer components that can cooperate to work synergistically via chromatin interactions. Furthermore, we revealed that combinational usage of GSK343 and X5050 could potentially lead to cancer ablation through disruption of “super-silencers”.

## Methods

We performed Hi-C, ChIP-seq, RNA-seq, HiChIP, cell culture, RT-qPCR, CRISPR excision, 4C-seq, 3C-PCR, xenograft models, western blot, cell cycle analysis, colony formation assay, adhesion assays, siRNA knock down experiment and growth curves as described in the **Supplementary Methods**. A list of all libraries used and generated is provided in **Supplementary Data 6**. A list of all the primers used is provided in **Supplementary Table S1**.

## Acknowledgements

This research is supported by a National Research Foundation Competitive Research Programme grant awarded to V.T. as lead PI and M.J.F. as co-PI (NRF-CRP17-2017-02). This research is supported by the National Research Foundation Singapore and the Singapore Ministry of Education under its Research Centres of Excellence initiative. This research is supported by a Ministry of Education Tier II grant awarded to M.J.F (T2EP30120-0020).

## Author contributions

Y.Z., K.J.C., Y.C.C. and M.J.F. conceived of the research and contributed to the study design. Y.C.C. and K.J.C. performed bioinformatics analysis. Y.Z. designed and performed CRISPR knock out experiments, Hi-C, 4C, 3C-PCR, RNA-seq, ChIP-seq, ChIP-qPCR and other functional experiments for KO clones. Y.Z. designed and performed GSK343 and X5050 drug treatment experiments including drug treatment, RT-qPCR, RNA-seq and Hi-C. A.N. and M.O. performed the drug synergy, colony formation assay, cell cycle analysis and mice experiments for the drug treatment. A.R., L.M. and V.T. designed and performed the xenograft experiments for DKO. Y.X.S. helped with the GSK343 treated Hi-C preparation. C.Y.F. helped with the microscope imaging and analysis. Y.Z., K.J.C., Y.C.C. and M.J.F. reviewed the data and wrote the manuscript. All authors reviewed and approved the manuscript.

## Data deposition

The list of libraries used in the study is provided in **Supplementary Data 6**. All datasets have been deposited into GEO under accession number GSE193489. Private access token **apepeyskxbedrsz** can be used to view the private data.

## Author information

The authors declare that they have no competing interests.

Correspondence and requests for materials should be addressed to.

## References

Akincilar, S. C., Khattar, E., Boon, P. L., Unal, B., Fullwood, M. J., & Tergaonkar, V. (2016). Long-Range Chromatin Interactions Drive Mutant TERT Promoter Activation. Cancer Discov, 6(11), 1276–1291. doi:10.1158/2159-8290.CD-16-0177

Babu, D., & Fullwood, M. J. (2015). 3D genome organization in health and disease: emerging opportunities in cancer translational medicine. Nucleus, 6(5), 382–393. doi:10.1080/19491034.2015.1106676

Bannister, A. J., & Kouzarides, T. (2011). Regulation of chromatin by histone modifications. Cell Res, 21(3), 381–395. doi:10.1038/cr.2011.22

Boija, A., Klein, I. A., Sabari, B. R., Dall’Agnese, A., Coffey, E. L., Zamudio, A. V., … Young, R. A. (2018). Transcription Factors Activate Genes through the Phase-Separation Capacity of Their Activation Domains. Cell, 175(7), 1842–1855 e1816. doi:10.1016/j.cell.2018.10.042

Bradner, J. E., Hnisz, D., & Young, R. A. (2017). Transcriptional Addiction in Cancer. Cell, 168(4), 629–643. doi:10.1016/j.cell.2016.12.013

Cai, Y., Zhang, Y., Loh, Y. P., Tng, J. Q., Lim, M. C., Cao, Z., … Fullwood, M. J. (2021). H3K27me3-rich genomic regions can function as silencers to repress gene expression via chromatin interactions. Nat Commun, 12(1), 719. doi:10.1038/s41467-021-20940-y

Cao, F., Fang, Y., Tan, H. K., Goh, Y., Choy, J. Y. H., Koh, B. T. H., … Fullwood, M. J. (2017). Super-Enhancers and Broad H3K4me3 Domains Form Complex Gene Regulatory Circuits Involving Chromatin Interactions. Sci Rep, 7(1), 2186. doi:10.1038/s41598-017-02257-3

Charbord, J., Poydenot, P., Bonnefond, C., Feyeux, M., Casagrande, F., Brinon, B., … Perrier, A. L. (2013). High throughput screening for inhibitors of REST in neural derivatives of human embryonic stem cells reveals a chemical compound that promotes expression of neuronal genes. Stem Cells, 31(9), 1816–1828. doi:10.1002/stem.1430

Chiabrando, D., Vinchi, F., Fiorito, V., Mercurio, S., & Tolosano, E. (2014). Heme in pathophysiology: a matter of scavenging, metabolism and trafficking across cell membranes. Front Pharmacol, 5, 61. doi:10.3389/fphar.2014.00061

Deng, W., Lee, J., Wang, H., Miller, J., Reik, A., Gregory, P. D., … Blobel, G. A. (2012). Controlling long-range genomic interactions at a native locus by targeted tethering of a looping factor. Cell, 149(6), 1233–1244. doi:10.1016/j.cell.2012.03.051

Doni Jayavelu, N., Jajodia, A., Mishra, A., & Hawkins, R. D. (2020). Candidate silencer elements for the human and mouse genomes. Nat Commun, 11(1), 1061. doi:10.1038/s41467-020-14853-5

Flavahan, W. A., Drier, Y., Liau, B. B., Gillespie, S. M., Venteicher, A. S., Stemmer-Rachamimov, A. O., … Bernstein, B. E. (2016). Insulator dysfunction and oncogene activation in IDH mutant gliomas. Nature, 529(7584), 110–114. doi:10.1038/nature16490

Frankel, N., Davis, G. K., Vargas, D., Wang, S., Payre, F., & Stern, D. L. (2010). Phenotypic robustness conferred by apparently redundant transcriptional enhancers. Nature, 466(7305), 490–493. doi:10.1038/nature09158

Gasparian, A. V., Burkhart, C. A., Purmal, A. A., Brodsky, L., Pal, M., Saranadasa, M., … Gurova, K. V. (2011). Curaxins: anticancer compounds that simultaneously suppress NF-kappaB and activate p53 by targeting FACT. Sci Transl Med, 3(95), 95ra74. doi:10.1126/scitranslmed.3002530

Hay, D., Hughes, J. R., Babbs, C., Davies, J. O. J., Graham, B. J., Hanssen, L., … Higgs, D. R. (2016). Genetic dissection of the alpha-globin super-enhancer in vivo. Nat Genet, 48(8), 895–903. doi:10.1038/ng.3605

Hietakangas, V., Poukkula, M., Heiskanen, K. M., Karvinen, J. T., Sistonen, L., & Eriksson, J. E. (2003). Erythroid differentiation sensitizes K562 leukemia cells to TRAIL-induced apoptosis by downregulation of c-FLIP. Mol Cell Biol, 23(4), 1278–1291. doi:10.1128/MCB.23.4.1278-1291.2003

Hnisz, D., Abraham, B. J., Lee, T. I., Lau, A., Saint-Andre, V., Sigova, A. A., … Young, R. A. (2013). Super-enhancers in the control of cell identity and disease. Cell, 155(4), 934–947. doi:10.1016/j.cell.2013.09.053

Hong, J. W., Hendrix, D. A., & Levine, M. S. (2008). Shadow enhancers as a source of evolutionary novelty. Science, 321(5894), 1314. doi:10.1126/science.1160631

Huang, D., Petrykowska, H. M., Miller, B. F., Elnitski, L., & Ovcharenko, I. (2019). Identification of human silencers by correlating cross-tissue epigenetic profiles and gene expression. Genome Res, 29(4), 657–667. doi:10.1101/gr.247007.118

Huang, J., Li, K., Cai, W., Liu, X., Zhang, Y., Orkin, S. H., … Yuan, G. C. (2018). Dissecting super-enhancer hierarchy based on chromatin interactions. Nat Commun, 9(1), 943. doi:10.1038/s41467-018-03279-9

Huang, J., Marco, E., Pinello, L., & Yuan, G. C. (2015). Predicting chromatin organization using histone marks. Genome Biol, 16, 162. doi:10.1186/s13059-015-0740-z

Hwang, J. Y., & Zukin, R. S. (2018). REST, a master transcriptional regulator in neurodegenerative disease. Curr Opin Neurobiol, 48, 193–200. doi:10.1016/j.conb.2017.12.008

Kantidze, O. L., Luzhin, A. V., Nizovtseva, E. V., Safina, A., Valieva, M. E., Golov, A. K., … Razin, S. V. (2019). The anti-cancer drugs curaxins target spatial genome organization. Nat Commun, 10(1), 1441. doi:10.1038/s41467-019-09500-7

Karimian, A., Ahmadi, Y., & Yousefi, B. (2016). Multiple functions of p21 in cell cycle, apoptosis and transcriptional regulation after DNA damage. DNA Repair (Amst), 42, 63–71. doi:10.1016/j.dnarep.2016.04.008

Karlic, R., Chung, H. R., Lasserre, J., Vlahovicek, K., & Vingron, M. (2010). Histone modification levels are predictive for gene expression. Proc Natl Acad Sci U S A, 107(7), 2926–2931. doi:10.1073/pnas.0909344107

Kaufmann, S. H., Desnoyers, S., Ottaviano, Y., Davidson, N. E., & Poirier, G. G. (1993). Specific proteolytic cleavage of poly(ADP-ribose) polymerase: an early marker of chemotherapy-induced apoptosis. Cancer Res, 53(17), 3976–3985.

Kim, S., & Shendure, J. (2019). Mechanisms of Interplay between Transcription Factors and the 3D Genome. Mol Cell, 76(2), 306–319. doi:10.1016/j.molcel.2019.08.010

Larson, A. G., Elnatan, D., Keenen, M. M., Trnka, M. J., Johnston, J. B., Burlingame, A. L., … Narlikar, G. J. (2017). Liquid droplet formation by HP1alpha suggests a role for phase separation in heterochromatin. Nature, 547(7662), 236–240. doi:10.1038/nature22822

Li, L., Lyu, X., Hou, C., Takenaka, N., Nguyen, H. Q., Ong, C. T., … Corces, V. G. (2015). Widespread rearrangement of 3D chromatin organization underlies polycomb-mediated stress-induced silencing. Mol Cell, 58(2), 216–231. doi:10.1016/j.molcel.2015.02.023

Li, Q., Barkess, G., & Qian, H. (2006). Chromatin looping and the probability of transcription. Trends Genet, 22(4), 197–202. doi:10.1016/j.tig.2006.02.004

Long, H. K., Prescott, S. L., & Wysocka, J. (2016). Ever-Changing Landscapes: Transcriptional Enhancers in Development and Evolution. Cell, 167(5), 1170–1187. doi:10.1016/j.cell.2016.09.018

Loven, J., Hoke, H. A., Lin, C. Y., Lau, A., Orlando, D. A., Vakoc, C. R., … Young, R. A. (2013). Selective inhibition of tumor oncogenes by disruption of super-enhancers. Cell, 153(2), 320–334. doi:10.1016/j.cell.2013.03.036

Lupianez, D. G., Kraft, K., Heinrich, V., Krawitz, P., Brancati, F., Klopocki, E., … Mundlos, S. (2015). Disruptions of topological chromatin domains cause pathogenic rewiring of gene-enhancer interactions. Cell, 161(5), 1012–1025. doi:10.1016/j.cell.2015.04.004

Ma, Y., Wang, L., Neitzel, L. R., Loganathan, S. N., Tang, N., Qin, L., … Wang, J. (2017). The MAPK Pathway Regulates Intrinsic Resistance to BET Inhibitors in Colorectal Cancer. Clin Cancer Res, 23(8), 2027–2037. doi:10.1158/1078-0432.CCR-16-0453

Ma, Y. N., Chen, M. T., Wu, Z. K., Zhao, H. L., Yu, H. C., Yu, J., & Zhang, J. W. (2013). Emodin can induce K562 cells to erythroid differentiation and improve the expression of globin genes. Mol Cell Biochem, 382(1-2), 127–136. doi:10.1007/s11010-013-1726-3

Mah, L. J., El-Osta, A., & Karagiannis, T. C. (2010). gammaH2AX: a sensitive molecular marker of DNA damage and repair. Leukemia, 24(4), 679–686. doi:10.1038/leu.2010.6

Martin, S. J., Bradley, J. G., & Cotter, T. G. (1990). HL-60 cells induced to differentiate towards neutrophils subsequently die via apoptosis. Clin Exp Immunol, 79(3), 448–453. doi:10.1111/j.1365-2249.1990.tb08110.x

Moorthy, S. D., Davidson, S., Shchuka, V. M., Singh, G., Malek-Gilani, N., Langroudi, L., … Mitchell, J. A. (2017). Enhancers and super-enhancers have an equivalent regulatory role in embryonic stem cells through regulation of single or multiple genes. Genome Res, 27(2), 246–258. doi:10.1101/gr.210930.116

Narendra, V., Bulajic, M., Dekker, J., Mazzoni, E. O., & Reinberg, D. (2016). CTCF-mediated topological boundaries during development foster appropriate gene regulation. Genes Dev, 30(24), 2657–2662. doi:10.1101/gad.288324.116

Ngan, C. Y., Wong, C. H., Tjong, H., Wang, W., Goldfeder, R. L., Choi, C., … Wei, C. L. (2020). Chromatin interaction analyses elucidate the roles of PRC2-bound silencers in mouse development. Nat Genet, 52(3), 264–272. doi:10.1038/s41588-020-0581-x

Nora, E. P., Goloborodko, A., Valton, A. L., Gibcus, J. H., Uebersohn, A., Abdennur, N., … Bruneau, B. G. (2017). Targeted Degradation of CTCF Decouples Local Insulation of Chromosome Domains from Genomic Compartmentalization. Cell, 169(5), 930–944 e922. doi:10.1016/j.cell.2017.05.004

Osterwalder, M., Barozzi, I., Tissieres, V., Fukuda-Yuzawa, Y., Mannion, B. J., Afzal, S. Y., … Pennacchio, L. A. (2018). Enhancer redundancy provides phenotypic robustness in mammalian development. Nature, 554(7691), 239–243. doi:10.1038/nature25461

Pennacchio, L. A., Bickmore, W., Dean, A., Nobrega, M. A., & Bejerano, G. (2013). Enhancers: five essential questions. Nat Rev Genet, 14(4), 288–295. doi:10.1038/nrg3458

Perino, M., & Veenstra, G. J. (2016). Chromatin Control of Developmental Dynamics and Plasticity. Dev Cell, 38(6), 610–620. doi:10.1016/j.devcel.2016.08.004

Portela, V. M., Dirandeh, E., Guerrero-Netro, H. M., Zamberlam, G., Barreta, M. H., Goetten, A. F., & Price, C. A. (2015). The role of fibroblast growth factor-18 in follicular atresia in cattle. Biol Reprod, 92(1), 14. doi:10.1095/biolreprod.114.121376

Pott, S., & Lieb, J. D. (2015). What are super-enhancers? Nat Genet, 47(1), 8–12. doi:10.1038/ng.3167

Rao, S. S., Huntley, M. H., Durand, N. C., Stamenova, E. K., Bochkov, I. D., Robinson, J. T., … Aiden, E. L. (2014). A 3D map of the human genome at kilobase resolution reveals principles of chromatin looping. Cell, 159(7), 1665–1680. doi:10.1016/j.cell.2014.11.021

Rowley, M. J., Nichols, M. H., Lyu, X., Ando-Kuri, M., Rivera, I. S. M., Hermetz, K., … Corces, V. G. (2017). Evolutionarily Conserved Principles Predict 3D Chromatin Organization. Mol Cell, 67(5), 837–852 e837. doi:10.1016/j.molcel.2017.07.022

Sabari, B. R., Dall’Agnese, A., Boija, A., Klein, I. A., Coffey, E. L., Shrinivas, K., … Young, R. A. (2018). Coactivator condensation at super-enhancers links phase separation and gene control. Science, 361(6400). doi:10.1126/science.aar3958

Shin, H. Y., Willi, M., HyunYoo, K., Zeng, X., Wang, C., Metser, G., & Hennighausen, L. (2016). Hierarchy within the mammary STAT5-driven Wap super-enhancer. Nat Genet, 48(8), 904–911. doi:10.1038/ng.3606

Stock, J. K., Giadrossi, S., Casanova, M., Brookes, E., Vidal, M., Koseki, H., … Pombo, A. (2007). Ring1-mediated ubiquitination of H2A restrains poised RNA polymerase II at bivalent genes in mouse ES cells. Nat Cell Biol, 9(12), 1428–1435. doi:10.1038/ncb1663

Subramanian, A., Tamayo, P., Mootha, V. K., Mukherjee, S., Ebert, B. L., Gillette, M. A., … Mesirov, J. P. (2005). Gene set enrichment analysis: a knowledge-based approach for interpreting genome-wide expression profiles. Proc Natl Acad Sci U S A, 102(43), 15545–15550. doi:10.1073/pnas.0506580102

Thongjuea, S., Stadhouders, R., Grosveld, F. G., Soler, E., & Lenhard, B. (2013). r3Cseq: an R/Bioconductor package for the discovery of long-range genomic interactions from chromosome conformation capture and next-generation sequencing data. Nucleic Acids Res, 41(13), e132. doi:10.1093/nar/gkt373

van Mierlo, G., Veenstra, G. J. C., Vermeulen, M., & Marks, H. (2019). The Complexity of PRC2 Subcomplexes. Trends Cell Biol, 29(8), 660–671. doi:10.1016/j.tcb.2019.05.004

van Steensel, B., & Furlong, E. E. M. (2019). The role of transcription in shaping the spatial organization of the genome. Nat Rev Mol Cell Biol, 20(6), 327–337. doi:10.1038/s41580-019-0114-6

Visel, A., Blow, M. J., Li, Z., Zhang, T., Akiyama, J. A., Holt, A., … Pennacchio, L. A. (2009). ChIP-seq accurately predicts tissue-specific activity of enhancers. Nature, 457(7231), 854–858. doi:10.1038/nature07730

Werner, T., Hammer, A., Wahlbuhl, M., Bosl, M. R., & Wegner, M. (2007). Multiple conserved regulatory elements with overlapping functions determine Sox10 expression in mouse embryogenesis. Nucleic Acids Res, 35(19), 6526–6538. doi:10.1093/nar/gkm727

Whyte, W. A., Orlando, D. A., Hnisz, D., Abraham, B. J., Lin, C. Y., Kagey, M. H., … Young, R. A. (2013). Master transcription factors and mediator establish super-enhancers at key cell identity genes. Cell, 153(2), 307–319. doi:10.1016/j.cell.2013.03.035

Wutz, G., Varnai, C., Nagasaka, K., Cisneros, D. A., Stocsits, R. R., Tang, W., … Peters, J. M. (2017). Topologically associating domains and chromatin loops depend on cohesin and are regulated by CTCF, WAPL, and PDS5 proteins. EMBO J, 36(24), 3573–3599. doi:10.15252/embj.201798004

Zhang, T., Cooper, S., & Brockdorff, N. (2015). The interplay of histone modifications - writers that read. EMBO Rep, 16(11), 1467–1481. doi:10.15252/embr.201540945

